# Prediction of treatment response in infantile epileptic spasms syndrome using EEG phase-amplitude coupling

**DOI:** 10.1101/2025.09.19.677383

**Authors:** S. Mostaghimi, Marco A. Pinto-Orellana, Nathaniel Green, Daniel W. Shrey, Makoto Miyakoshi, Shaun A. Hussain, Beth A. Lopour

## Abstract

**Objective:** Treatment selection for infantile epileptic spasms syndrome (IESS) is complex and multifaceted, and currently no EEG biomarkers can guide this decision by predicting treatment response. We tested the predictive value of phase-amplitude coupling (PAC), as IESS patients are known to have elevated PAC.

**Methods:** We analyzed retrospective EEG recordings from 40 IESS patients, before and after treatment, and 20 healthy controls. Patients were classified as responders (n=25) or non-responders (n=15) based on short-term treatment outcomes. We measured PAC in each EEG using modulation index (MI) and mean vector length (MVL) and analyzed the relationship between pre- and post-treatment values and the ability of pre-treatment values to predict response.

**Results:** MI and MVL values decreased with treatment in almost all subjects. However, non-responders had significantly higher pre-treatment MI than responders (P<0.05), suggesting utility for predicting treatment response. Logistic regression modeling suggested that a 0.5 unit decrease in log(MI), which is approximately one IQR of the pre-treatment log(MI) values, results in a 6-fold increase in odds of positive treatment response.

**Significance:** MI reflects short-term treatment response and is a candidate predictive EEG biomarker for IESS. MI may offer individualized insights for treatment selection and management strategies for IESS.

**Key Points Box:**

- Infantile epileptic spasms syndrome (IESS) is associated with high phase-amplitude coupling (PAC) that decreases with treatment
- Low pre-treatment values of EEG PAC are predictive of positive short-term treatment response for IESS
- The results were strongest when PAC was measured using the modulation index, as this method is known to be robust to noise

## Introduction

Infantile epileptic spasms syndrome (IESS) is an epileptic encephalopathy that causes progressive cognitive decline and behavioral disabilities (Wong et al., 2001, Auvin et al., 2012), and long-term outcomes are worse if the disease is not diagnosed and treated promptly (O’Callaghan et al., 2011, Auvin et al., 2012, Hussain, 2018). First-line treatments for IESS include adrenocorticotropic hormone (ACTH), corticosteroids (prednisone, prednisolone), vigabatrin, or the combination of hormonal therapy (ACTH or prednisolone) and vigabatrin (Velíšek et al., 2020). Limited data suggest that the ketogenic diet might be a viable first-line therapy (Dressler et al., 2019), and a small fraction of IESS patients with drug-resistant epileptic spasms are considered for surgical treatment (Hussain, 2018). Beyond comparisons of efficacy, the selection of specific treatments is complicated by varied risks and costs. Whereas hormonal therapy carries the risk of immunosuppression and potentially lethal infection (Hussain, 2018), vigabatrin has been linked to irreversible peripheral visual field defects (Gaily et al., 2009, Riikonen, 2020). Knowledge of a patient’s likelihood of responding to pharmaceutical treatment could guide patient management and help avoid unnecessary treatments with risks of severe side effects.

Computational EEG metrics have been proposed as biomarkers relevant to the diagnosis and treatment of IESS. Studies have used computational analysis to quantify differences in EEG features between IESS and control patients, such as amplitude, spectral power (Burroughs et al., 2014, Smith et al., 2018, Smith et al., 2021), entropy (Chu et al., 2021, Smith et al., 2021), and functional connectivity (Shrey et al., 2018, Stacey et al., 2020, Smith et al., 2021, Romero Mila et al., 2022). Such biomarkers might also be helpful for assessing IESS treatment response, for example, EEG complexity (entropy) (Chu et al., 2021, Zhang et al., 2023), relative power spectrum (Kanai et al., 2022), functional connectivity (Shrey et al., 2018, Kanai et al., 2022, Kim et al., 2023), and long-range temporal correlations (Smith et al., 2017, Rajaraman et al., 2024). The Burden of AmplitudeS and Epileptiform Discharges (BASED) score (Mytinger et al., 2021) is a metric derived from visual scoring of the EEG. The results associated with this metric have been mixed. Some studies suggest that the BASED score can predict short-term treatment response and aid in identifying IESS patients and measuring their treatment responses (Mytinger et al., 2021, Wan et al., 2022). However, other studies have shown that the BASED score fails to reliably predict short-term treatment outcomes (Rajaraman et al., 2024).

One computational metric that has shown particular promise as a biomarker of epilepsy is phase-amplitude Coupling (PAC), which measures the modulation of the EEG amplitude at high frequencies by the phase of lower frequencies (Tort et al., 2010). While there are many methods for assessing PAC, the Mean Vector Length (MVL) (Canolty et al., 2006) and Modulation Index (MI) (Tort et al., 2010) are two of the most commonly used. For instance, the value of MI increases during preictal periods relative to interictal periods; it has thus been suggested as a potential tool for seizure prediction and detection (Jiang et al., 2023). Moreover, PAC has facilitated the localization of epileptogenic tissue for various types of epilepsy, with higher MVL values in these regions (Iimura et al., 2022, Wada et al., 2022). Specifically for IESS, MVL values for coupling between the delta- and gamma-bands and the delta- and high-frequency oscillation (HFO) bands were higher in IESS compared to controls (Nariai et al., 2020, Miyakoshi et al., 2021). In IESS patients who underwent multilobe resection and achieved seizure freedom, the resected brain regions exhibited strong coupling between the delta-band and fast-ripple bands (Iimura et al., 2018). Another study that investigated post-surgical changes in PAC showed that delta-gamma MVL values significantly decreased in non-lesional IESS patients who underwent disconnection surgeries and achieved seizure freedom (Uda et al., 2021). Only one study has examined PAC as a biomarker of IESS treatment response to hormonal therapies; they found no differences in coupling strengths between responders and non-responders (0.5-8 Hz low-frequency band, coupled with ripples or fast ripples) but suggested that the preferred phase angles differed (Bernardo et al., 2020). A systematic examination of PAC in IESS patients before and after treatment with hormonal therapies, as well as a comparison to control values, has yet to be completed.

Therefore, we evaluated MI and MVL as biomarkers for the assessment and prediction of treatment response in IESS, with healthy controls as an approximately age-matched comparison group. Based on prior studies, we hypothesized: 1) IESS treatment will lead to a normalization of PAC values, such that they are more similar to those of control patients, and 2) the change in PAC value due to treatment will reflect treatment response, with lower values reflecting successful treatment.

## 1 Methods

### 1.1 Subject Recruitment

Approval to conduct this retrospective study was obtained from the Institutional Review Board at the University of California, Los Angeles. We analyzed EEG data from subjects with IESS and healthy controls; details on the subject selection and data collection can be found in (Miyakoshi et al., 2021, Smith et al., 2021). Briefly, 50 subjects with IESS (cases) were identified by evaluation of their overnight video-EEG recording for epileptic spasms, regardless of the presence or absence of hypsarrhythmia. Another 50 subjects underwent the same evaluation, but they did not exhibit any seizures and were determined to be neurologically normal (controls). To create approximately age-matched groups, a random selection process was used to select 20 controls (median age 11.0 months, IQR 7.6–22.6) and 40 cases (median age 9.9 months, IQR 7.4–27.9). For this study, we analyzed EEG recordings from these subsets of cases and controls.

For the cases, EEGs were collected from two overnight video-EEG sessions. The first session, termed “pre-treatment,” involved collecting baseline EEG recordings before patients began treatment. Following this initial recording, patients received their prescribed therapies (Table 1). Approximately two weeks later (median interval of 16 days, IQR 12-20), a “post-treatment” EEG recording session was conducted.

**Table 1.** Demographics and interventions used for IESS patients.

|  |  | Aged-matched cases<br>(n=40) |
| --- | --- | --- |
| Age at pre-treatment EEG, months<sup>1</sup> |  | 11.0 [7.6, 22.6] |
| Age at onset of epileptic spasms, months<sup>1</sup> |  | 7.4 [4.6, 11.8] |
| Female, n (%) |  | 20 (50%) |
| Treatment<br>after the<br>baseline<br>EEG | Prednisone, n (%) | 26 (65%) |
|  | Vigabatrin, n (%) | 9 (22.5%) |
|  | Adrenocorticotrophic (ACTH),<br>n (%) | 5 (12.5%) |

<sup>1</sup>Median [interquartile range].

Responders were defined as cases that had normalization of the EEG and clinically assessed resolution of epileptic spams at 28 days from the start of the treatment (n=25). Cases that exhibited continued spasms and/or did not have normalization of EEG after treatment were designated as non-responders (n=15).

### 1.2 Electroencephalogram (EEG) recordings

Overnight scalp EEG recordings were captured using Nihon Kohden EEG acquisition systems (Irvine, California). Twenty-one gold-plated electrodes were placed following the international 10-20 system, which included 19 EEG channels (Fp1, Fp2, F3, F4, C3, C4, P3, P4, O1, O2, F7, F8, T3, T4, T5, T6, Fz, Cz, and Pz) and two electrodes placed at the mastoids. The recordings were digitally sampled at either 200 Hz or 2000 Hz. EEG recordings that had been sampled at 2000 Hz were subsequently resampled to 200 Hz before undergoing further preprocessing.

EEG clip selection for each patient is explained in detail in (Smith et al., 2021). In summary, for each overnight EEG, two sleep EEG clips were extracted from each patient (median clip duration 27.5 minutes, IQR 19.9-29.9). Selection of each clip began at a randomly selected time point in the EEG; a reviewer then scanned forward in time until the subject was determined to be sleeping, and the clip was extracted.

### 1.3 Preprocessing and artifact detection

All EEG recordings were re-referenced to the common average, then bandpass filtered in the delta (3-4 Hz) and gamma (35-70 Hz) frequency bands. These frequency bands were chosen to match a previous study that demonstrated differences in PAC between IESS patients and controls (Miyakoshi et al., 2021). Digital filters were implemented using the specifications described in (Miyakoshi et al., 2021), with the addition of a second-order infinite impulse response notch filter at 60 Hz, with a bandwidth of 0.01 Hz, for the removal of electrical line interference.

EEG artifacts were identified using the methods described in (Durka et al., 2003, Moretti et al., 2003, Smith et al., 2021), where the segments with very high z-scored amplitude in any channel were defined as artifactual. Subsequently, both EEG clips for all patients were divided into 6-second epochs. Epochs containing artifacts were excluded from the analysis for all channels.

After these preprocessing steps, the smallest number of clean epochs across all subjects and both EEG recordings was determined to be n=79. Therefore, to equalize the amount of data across subjects, the first 79 clean epochs for each patient and each EEG were selected for analysis, and any additional clean epochs were excluded.

### 1.4 Phase-amplitude coupling analysis

PAC is a type of cross-frequency coupling in which the signal phase in a low-frequency band (phase-providing frequency) is related to the signal amplitude in a higher frequency band (amplitude-providing frequency). Here we measure PAC using two metrics, MI (Tort et al., 2010) and MVL (Canolty et al., 2006). MVL was used in previous studies investigating PAC in IESS, and its numerical value is directly proportional to the amplitude of the amplitude-providing frequency band. MI has been shown to be more robust to noise than MVL, and its value is also not directly affected by the amplitude of the amplitude-providing frequency band (Hülsemann et al., 2019). Therefore, MI and MVL exhibit complementary strengths, making it advantageous to use both in our study.

We considered the delta-band signal to be the phase-contributing component (*f*_*P*_), and the gamma-band signal to be the amplitude-modulated component (*f*_*A*_), where *f* stands for the filtered signal and subscripts A and P refer to the amplitude- and phase-providing frequencies, respectively. Using the Hilbert transform, we calculated the instantaneous amplitude of *f*_*A*_, denoted by *X*_*A*_(*t*), and the instantaneous phase of *f*_*P*_, denoted by *θ*_*P*_(*t*). Then MVL was defined as follows, where N is the total number of data points in the 6-second epoch:

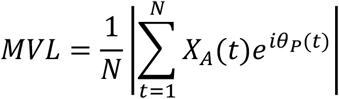

To calculate MI, we used a joint bivariate time series of the delta phase and the gamma amplitude defined as (*θ*_*P*_(*t*), *X*_*A*_(*t*)). Then, we binned *θ*_*P*_(*t*) into M=18 bins of 20 degrees and calculated the average gamma-band amplitude associated with each phase bin j, denoted by 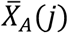. Next, we normalized each 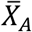 by the sum of all averages over all M bins:

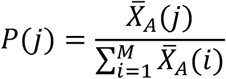

Due to this normalization, P(j) is also known as a “pseudo-probability” because the values of P(j) sum to 1. Then, we calculated the Kullback–Leibler (KL) divergence of P(j) from a uniform distribution U(j) using the following equation:

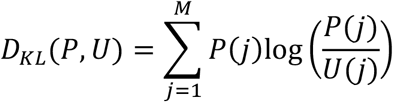

Finally, MI is obtained as follows:

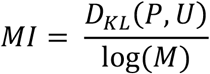

We calculated the MI and MVL values of the first 79 epochs of each sleep EEG clip in all 19 channels for every patient. We then took the median across all 158 epochs from the two sleep EEG clips for MI and MVL for each electrode in every patient.

### 1.5 Logistic regression

We employed logistic regression to assess whether PAC could aid in predicting short-term response to IESS treatment. We also included lead time in our model, as a previous study showed that lead time is associated with treatment response (Rajaraman et al., 2024). We followed their approach of utilizing an approximate log-transform of the lead time and converting it into a discrete ordinal variable “LEADTIME” such that 0 represents a lead time of 0 – 7 days, 1 represents 8 – 14 days, 2 represents 15 – 30 days, 3 represents 31 – 60 days, and 4 represents a lead time greater than 60 days (O’Callaghan et al., 2011).

Subsequently, we used a binomial logistic regression model with treatment response as the dependent variable and MI, MVL, and LEADTIME as the independent variables, either alone or in combination. We categorized treatment responses as “responder” (1) or “non-responder” (0). To facilitate a meaningful comparison between the models based on MI, MVL, and LEADTIME we log-transformed the PAC values for the pre-treatment cases. Then we computed the median PAC value across all 158 epochs of EEG to obtain one data point for each channel in each patient. Finally, to have a single data point for each patient, we took the median across all 19 channels.

### 1.6 Statistical analysis

We used the Wilcoxon signed-rank test to compare the paired median values of MI or MVL in pre- and post-treatment subjects for both responders and non-responders. Specifically, we compared all pre-treatment to all post-treatment cases, pre-treatment responders to post-treatment responders, and pre-treatment non-responders to post-treatment non-responders.

We performed the Mann-Whitney U-test for the comparison of MI or MVL between groups with unpaired data. Specifically, we compared pre-treatment cases to controls, post-treatment cases to controls, pre-treatment responders to pre-treatment non-responders, and post-treatment responders to post-treatment non-responders.

For all tests, we used the borderline significance level of p<0.05 and false discovery rate (FDR)-corrected p-value <0.05 to identify channels that showed statistical significance.

## 2 Results

### 2.1 Phase-amplitude coupling is higher in IESS

We first compared the PAC values of controls to pre-treatment cases and controls to post-treatment cases. Both pre- and post-treatment cases exhibited higher MI values compared to control subjects in all channels (Figure 1A). For MVL, none of the pre-treatment values were significantly different from controls (Figure 1B). In contrast, post-treatment cases exhibited statistically lower values than control subjects in all channels (Figure 1B). These trends were further confirmed when visualizing the median MI values (Figure 2A) and MVL values (Figure 2B), with the median taken across all epochs and then all channels in each subject.

**Figure 1.**
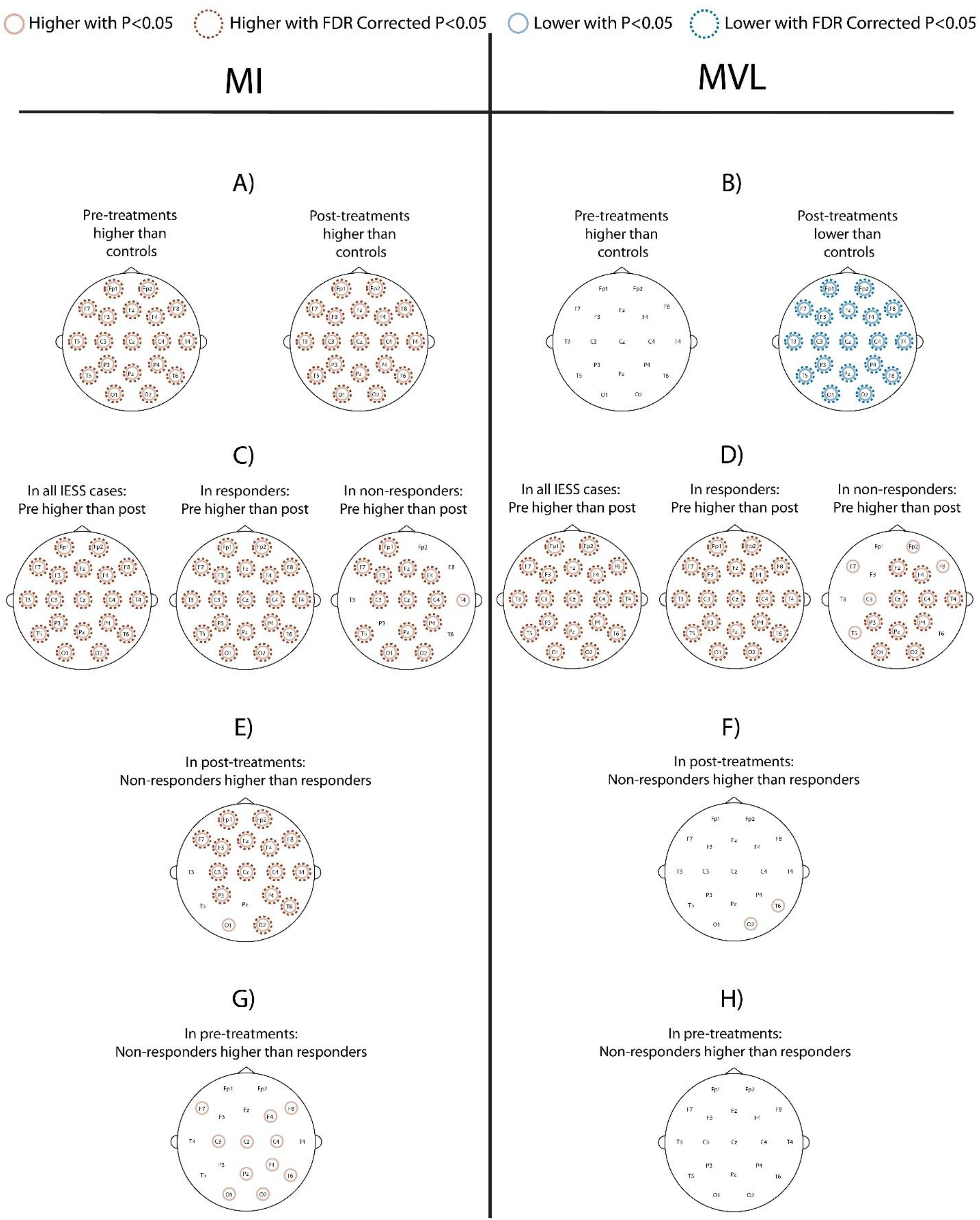
Head maps for MI and MVL displaying channels with statistical significance for the comparison in the subfigure title. Channels that were statistically significant with p<0.05 are marked with solid circles. The channels that maintained significance with FDR correction are additionally marked with dashed circles. Abbreviations: MI = modulation index, MVL = mean vector length, FDR = false discovery rate

**Figure 2.**
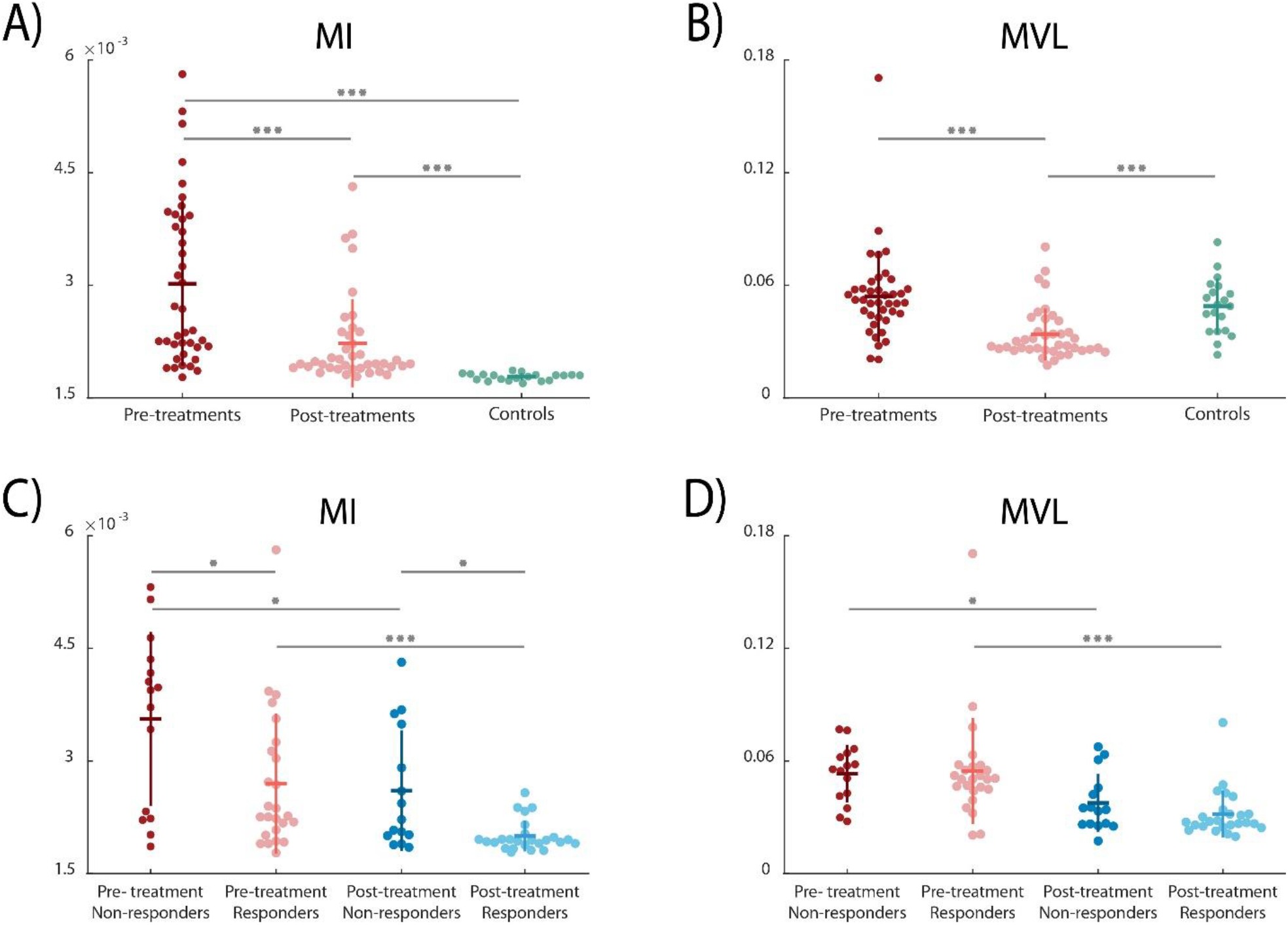
Beeswarm plots of median PAC in controls and subpopulations of IESS patients. For each group, the horizontal line represents the average of the data, while the vertical line indicates one standard deviation above and below the average of the data. The data shown are (A) median MI and (B) median MVL values, taken across all epochs and all channels, for controls and pre- and post-treatment IESS cases. (C) Median MI and (D) median MVL for IESS cases, subdivided into those that were ultimately responders or non-responders. Comparisons connected by horizontal lines indicate statistically significant differences. Asterisks denote the level of significance: ^*^ p < 0.05, ^**^ p < 0.01, and ^***^ p < Abbreviations: PAC = phase-amplitude coupling, IESS = infantile epileptic spasms syndrome, MI = modulation index, MVL = mean vector length

### 2.2 Phase-amplitude coupling decreases with treatment

Next, we compared the PAC values of the pre-treatment and post-treatment groups. Pre-treatment patients had significantly higher MI and MVL than post-treatment patients in all channels (Figure 1C, D, Figure 2A, B). This was also true when the responder and non-responder groups were tested independently, with almost all channels showing significantly higher PAC values for pre-treatment compared to post-treatment (Figure 1C, D, Figure 2C, D). This decrease due to treatment was consistent across subjects, as indicated by each patient’s median pre-treatment PAC value relative to their post-treatment PAC value (Figure 3).

**Figure 3.**
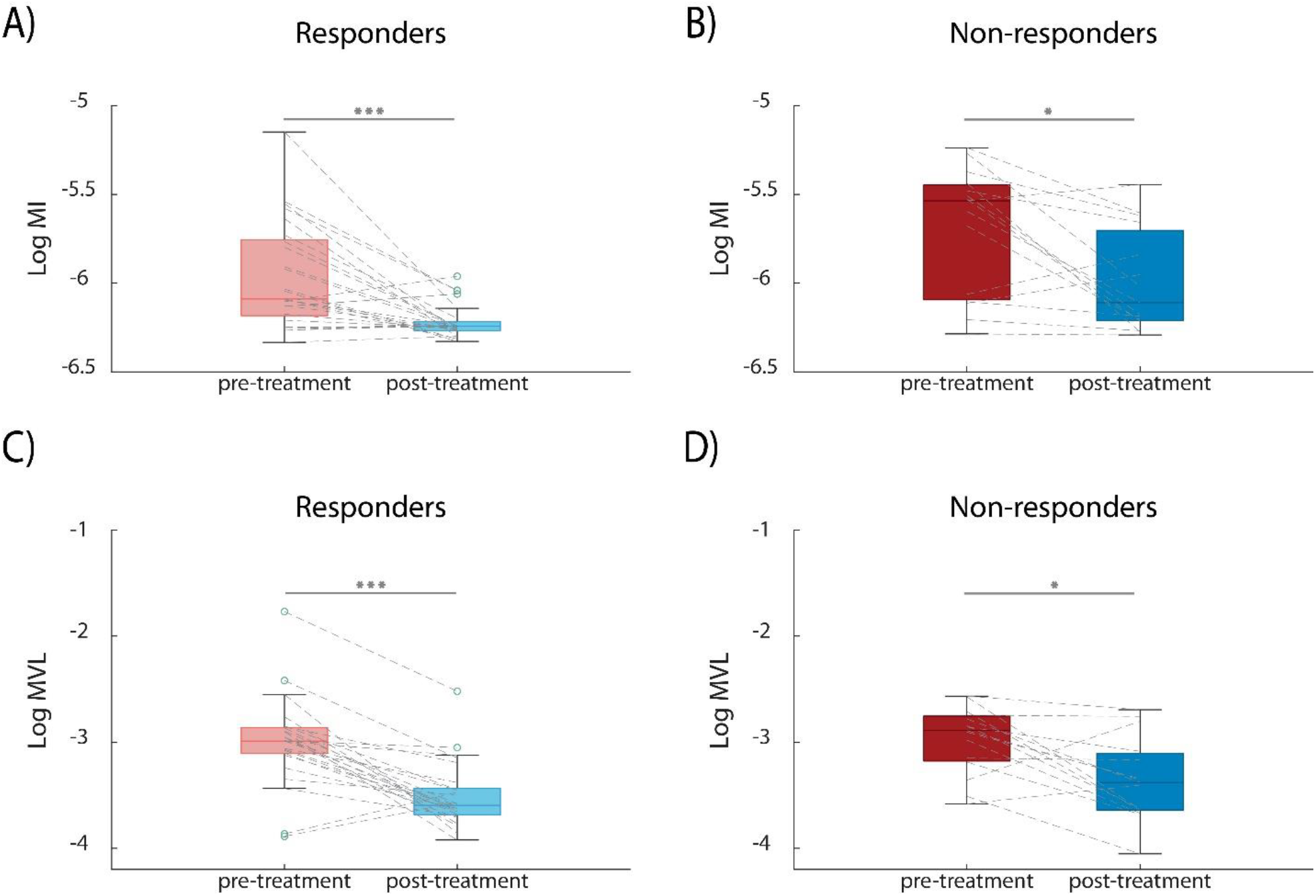
PAC decreases after IESS treatment. Subfigures show the log-scaled pre- and post-treatment values of PAC for A) MI in responders, B) MI in non-responders, C) MVL in responders, and D) MVL in non-responders. The value for each patient was obtained by taking the median over all 158 epochs and 19 channels. Green circles indicate outliers. Comparisons connected by horizontal lines indicate statistically significant differences. Asterisks denote the level of significance: ^*^ p < 0.05, ^**^ p < 0.01, and ^***^ p < 0.001. Abbreviations: PAC = phase-amplitude coupling, IESS = infantile epileptic spasms syndrome, MI = modulation index, MVL = mean vector length

### 2.3 Post-treatment phase-amplitude coupling is higher in non-responders compared to responders

To understand the relationship between PAC and treatment response, we compared post-treatment non-responders to post-treatment responders. Post-treatment non-responders had higher MI than post-treatment responders (Figure 1E and Figure 2C). This difference was statistically significant in 16 of 19 EEG channels for MI. For MVL, only two channels (O2 and T6) showed statistically higher MVL in non-responders compared to responders, and these differences did not remain significant after FDR correction (Figure 1F). When visualizing the median value across all channels, post-treatment responders and non-responders had comparable values (Figure 2D).

### 2.4 MI may aid in prediction of IESS treatment response

#### 2.4.1 Statistical testing results

Finally, we tested whether pre-treatment PAC could predict short-term treatment response by comparing pre-treatment PAC values for those who would ultimately be responders to those who would be non-responders. Channels F4, C3, C4, P4, O1, O2, F7, F8, T6, Cz, and Pz had significantly higher MI in pre-treatment non-responders compared to pre-treatment responders (p<0.05, Figure 1G), and this trend was also evident in the median values of MI (Figure 2C). For MVL, none of the channels showed statistically higher MVL values in pre-treatment non-responders than in pre-treatment responders (Figure 1H). The median pre-treatment MVL values were comparable for responders and non-responders (Figure 2D).

#### 2.4.2 Logistic regression results

As illustrated in Table 2, sequential bivariate logistic regression analysis demonstrated that treatment response was associated with lower pre-treatment log(MI) and LEADTIME, but it was not associated with log(MVL). The odds ratio for log(MI) was 0.08, indicating that each unit decrease in pre-treatment log-scaled MI was associated with a 12-fold (i.e., 1/0.08) increased odds of treatment response. To put this in the context of our data, we can say that a 0.5 unit decrease in log(MI), which is approximately one IQR of the pre-treatment log(MI) values (Figure 3A,B), results in a 6-fold increase in odds of positive treatment response.

**Table 2.** Logistic regression results modeling patient responses to pharmaceutical therapy based on the lead time (LEADTIME), and/or Phase-Amplitude Coupling (PAC) measured using Mean Vector Length (MVL) and Modulation Index (MI). The table presents the coefficient, p-value (P), and odds ratio (OR) for each independent variable. The 97.5% and 2.5% confidence interval (CI) for coefficients and odds ratios is reported in parentheses.

| Dependent variable | Independent variable | PAC related variable |  |  | LEADTIME |  |  |
| --- | --- | --- | --- | --- | --- | --- | --- |
|  |  | Coefficient (95% CI) | P | OR (95% CI) | Coefficient (95% CI) | P | OR (95% CI) |
| Patient response to treatment | Pre-treatment log(MI) | -2.486 (-4.788, -0.48) | 0.022 | 0.08 (0.01,0.69) | - | - | - |
|  | Pre-treatment log(MVL) | -0.152 (-1.958, 1.61) | 0.86 | 0.86 (0.15,4.79) | - | - | - |
|  | LEADTIME | - | - | - | -0.527 (-0.215, -0.017) | 0.034 | 0.59 (0.36,0.96) |
|  | Pre-treatment log(MI) + LEADTIME | -2.156 (-4.506, -0.087) | 0.052 | 0.12 (0.01,1.02) | -0.462 (-1.046, -0.012) | 0.068 | 0.63 (0.38,1.03) |
|  | Pre-treatment log(MVL) + LEADTIME | -0.453 (-2.333, 1.279) | 0.61 | 0.64 (0.11,3.61) | -0.544 (-1.133, -0.104) | 0.031 | 0.58 (0.35,0.95) |

In multivariable logistic regression, the association of MI and treatment response appears to be independent of LEADTIME; both MI and LEADTIME were borderline statistically significant (p = 0.052 and p = 0.068, respectively). Note that we did not detect multicollinearity in the multivariable logistic regression model based on LEADTIME and PAC (variance inflation factor = 1.005 for MI, 1.019 for MVL).

## 3 Discussion

### 3.1 Summary

Here we have shown, for the first time, that EEG PAC reflects treatment response in patients with IESS (Figures 1-3), and it may also aid in the prediction of short-term treatment response (Table 2). These EEG changes were not localized to a particular brain region, as we saw significant results across the head (Figure 1). We also found that the MI metric of PAC produced more robust results than the MVL metric, consistent with the conclusions of Tort et al. (Tort et al., 2010).

### 3.2 MI is a candidate biomarker for IESS

Consistent with prior studies (Nariai et al., 2020, Miyakoshi et al., 2021), we demonstrated that PAC may serve as a biomarker of IESS by showing that MI is significantly higher in all EEG channels in pre-treatment IESS cases compared to controls. Moreover, both MI and MVL exhibit a decrease after treatment, which is more pronounced in responders, suggesting that they reflect treatment response. When we considered the post-treatment EEG, the fact that non-responders had higher MI and MVL than responders suggests that the treatment might have been less effective for non-responders. This underscores the value of PAC as a measure of treatment response. This interpretation gains support from our observation that, although there were some channels where non-responders showed significant differences between pre- and post-treatment values, MI and MVL were significantly higher across all electrode channels in responders. When comparing post-treatment IESS cases to controls, we observed lower MVL in the post-treatment cases, which contradicts our initial hypothesis. In contrast, MI showed statistically higher values in post-treatment cases compared to controls. This discrepancy could arise due to the direct influence of the gamma-band EEG amplitude on the measurement of MVL, as opposed to MI, which incorporates a normalization step for EEG amplitude (Tort et al., 2010).

### 3.3 MI is more tolerant to noise and may therefore be a more sensitive biomarker of IESS treatment response

We generally observed that MI yielded stronger results than MVL, with more channels showing statistical significance for MI when comparing cases to controls and responders to non-responders, both pre- and post-treatment. The lower number of statistically significant channels for MVL might explain the finding of Bernando et al., who observed higher MVL in pre-treatment non-responders compared to responders, but reported that the difference was not statistically significant (Bernardo et al., 2020). Consistent with that, our results from logistic regression showed that only MI was significant in predicting the short-term treatment response, with higher pre-treatment MI observed in non-responders compared to responders. This suggests that the initial condition of non-responders might have been different from responders, potentially requiring higher doses of medication or longer treatment courses to achieve similar outcomes as responders.

### 3.4 Methodological considerations

Several factors influenced our decision to select MI and MVL to quantify PAC. Our selection of MVL was chiefly guided by its implementation in previous studies, in which patients with IESS exhibited higher MVL than normal controls (Nariai et al., 2020, Miyakoshi et al., 2021). Regarding MI, multiple considerations guided our decision. First, simulations have shown that MI is more robust to noise than MVL (Hülsemann et al., 2019). Second, it is understood that while MI evaluates PAC with a more balanced sensitivity to both amplitude and phase values, MVL’s sensitivity leans heavily towards amplitude values (Tort et al., 2010, Hülsemann et al., 2019). Last, MVL is typically recommended for analyzing long data epochs recorded at high sampling rates and possessing a high signal-to-noise ratio. Conversely, MI is recommended for shorter epochs with lower sampling frequencies and reduced signal-to-noise ratio (Tort et al., 2010, Hülsemann et al., 2019), aligning well with our dataset’s characteristics.

As for the choice of frequency bands, prior studies of PAC in IESS patients have used the delta-band as the phase-contributing frequency band. While some of these studies focused on the coupling between the delta-band and HFO bands (ripples and fast ripples, > 80 Hz) to measure MVL (Iimura et al., 2018, Bernardo et al., 2020, Nariai et al., 2020), we opted to focus on delta-gamma coupling, consistent with (Miyakoshi et al., 2021). The 200 Hz sampling frequency of our EEG data precluded us from measuring coupling with the HFO bands. Moreover, it has been shown that both pathological HFOs and gamma-band activities in epileptic patients tend to couple with delta-band oscillations (Grigorovsky et al., 2020).

### 3.5 Limitations

Our study comes with some limitations. One of the constraints was that the EEGs in our dataset were collected on a single date following the initiation of treatment, which we used to evaluate the patient’s short-term response, and there was a range of time intervals between pre- and post-treatment EEG collection. Additionally, the treatment plans for IESS patients varied depending on the individual patient’s needs, and some had ongoing treatments at the time of the baseline EEG collection. These limitations are inherent to the retrospective nature of our study. In a prospective study, standardizing the treatment and interval between the baseline (pre-treatment) and follow-up (post-treatment) for each patient could reduce variability in our measurements. Furthermore, our data collection was confined to a single center and involved only 40 patients. To enhance the robustness and validate our findings, it is essential to expand our data collection to include multiple centers and a larger cohort of participants. The logistic regression analysis suggested that MI may have value for predicting treatment response, but the results could have been impacted by an insufficient number of observations or other confounding factors. A larger cohort of IESS patients would give us the power to assess the differences between responders and non-responders and the influence of confounding factors more accurately.

In this study, we analyzed EEG data collected during sleep, when artifacts are less prevalent, minimizing the likelihood that our measurements are affected by noise. However, despite our efforts to identify and remove as many artifacts as possible, it is feasible that some, such as muscle artifacts, may have remained in the data and impacted our estimates of PAC. It will also be vital in future studies to explore these PAC measurements during wakeful states, as was done in (Miyakoshi et al., 2021), to fully understand their dynamics across different consciousness levels and the relation to treatment response.

### 3.6 Conclusion

Here we show that MI is a candidate predictive marker for treatment response in IESS patients, suggesting new directions for future research and clinical approaches. Our results also highlight a limitation of MVL, despite its broader usage in related research. Overall, the identification of biomarkers for short-term treatment response in IESS has the potential to provide valuable insights into the future trajectory of treatment, aiding in treatment selection and indicating when continued intervention is necessary.

## 5 Acknowledgements

This work was supported in part by the John C. Hench Foundation and UCB Biopharma.

## 6 Declaration of Interests

None of the authors have potential conflicts of interest to be disclosed.

## Notes

**Declarations of Interest:** None of the authors have potential conflicts of interest to be disclosed.

### Competing Interest Statement

The authors have declared no competing interest.

